# Obesogenic diet impairs memory consolidation via the hippocampal endocannabinoid system

**DOI:** 10.1101/2025.07.04.663181

**Authors:** Eva-Gunnel Ducourneau, Yoottana Janthakhin, José F. Oliveira da Cruz, Julien Artinian, Serge Alfos, Jean-Christophe Helbling, Isabelle Matias, Ioannis Bakoyiannis, M Mateo N’Diaye, Clémentine Bosch-Bouju, Mylène Potier, Luigi Bellocchio, Arnau Busquets-Garcia, Pierre Trifilieff, Giovanni Marsicano, Guillaume Ferreira

**Affiliations:** Univ. Bordeaux, INRAE, Bordeaux INP, NutriNeuro, UMR 1286, Bordeaux, France; Univ. of Bordeaux, INSERM, NeuroCentre Magendie, U1215, Bordeaux, France

## Abstract

Although obesogenic high-fat/high-sugar diets impair memory function in humans and rodents, the underlying mechanisms remain elusive. Given that the brain endocannabinoid system and type-1 cannabinoid receptors (CB_1_R) control memory processes and are overactive under obesogenic conditions, we studied whether the effects of obesogenic diet consumption on memory function are dependent on this system. Using an object recognition memory (ORM) task in male mice, we showed that CB_1_R activity is required for obesogenic diet-induced impairment of long-term memory performance. This impairment was prevented by post-training systemic blockade of CB_1_R, which also normalized training-induced hippocampal cellular and synaptic overactivation. Consistently, obesogenic diet potentiated the increase of hippocampal endocannabinoid levels and enhanced CB_1_R expression induced by ORM, and genetic CB_1_R deletion from hippocampal glutamatergic neurons abolished diet-induced memory deficits. Strikingly, obesogenic diet enhanced the hippocampal mTOR pathway in a CB_1_R-dependent manner, and pharmacological mTOR inhibition after training rescued diet-induced ORM consolidation deficits. Together, these results establish how an obesogenic environment can lead to hippocampal overactivation of the endocannabinoid system and of the mTOR pathway to eventually impair memory consolidation. Thus, these results shed light on the mechanisms of diet-induced cognitive alterations and may pave the way to novel therapeutic strategies.

**Highlights:**

- Obesogenic diet induces long-term memory deficits, which are rescued by CB_1_R blockade
- CB_1_R blockade rescues diet-induced aberrant hippocampal activity/plasticity after training
- Obesogenic diet enhances hippocampal endocannabinoid levels and CB_1_R after training
- Deletion of hippocampal CB_1_R rescues diet-induced long-term memory deficits
- Obesogenic diet enhances hippocampal mTOR phosphorylation after training
- mTOR inhibition rescues diet-induced memory consolidation deficits

## Introduction

Obesity is one of the most serious public health challenges of the 21^st^ century, with a prevalence that has doubled since 1990 in adults, affecting 20% of the population worldwide. It is largely due to excessive intake of energy-dense foods, particularly high-fat and sugar diets (HFD; for review: [1]). In addition to being a risk factor for cardiometabolic diseases, a large fraction of overweight and obese patients shows cognitive dysfunctions. Clinical and experimental studies indicate that obesogenic environments favour learning and memory impairments that depend on the integrity of the hippocampus, such as episodic and spatial memory (for review: [2]). The prevalence of obesity in adolescents has risen dramatically in the past four decades and has been exacerbated by the COVID-19 pandemic [3]. Adolescents with obesity display blunted performance in hippocampal-dependent tasks such as geometric and visuospatial problems or relational and episodic memory [4–6] that can eventually significantly worsen academic performances (for review: [7]). This is worrisome since this developmental period is sensitive to environmental challenges such as food ([8]) and is a crucial period for hippocampal maturation ([9]), shaping cognitive functions for life. We and others demonstrated that obesogenic HFD consumption in rodents during adolescence has a more drastic impact on memory than similar exposure in adulthood, suggesting that this developmental period is particularly sensitive to HFD effects on memory (for review: [10]). In particular, we showed that 12 weeks of HFD intake starting at weaning and covering the peri-adolescent period, altered hippocampal activity and plasticity and impaired many forms of hippocampal-dependent long-term memory, including spatial, relational, contextual and object-based ones [11–16]. However, the cellular mechanisms underlying these peri-adolescent HFD consumption-induced cognitive impairments remain largely unclear.

Interestingly, the endocannabinoid (eCB) system is at the crossroads of obesity and memory functions (for review: [17]). Clinical and experimental studies indicate that the eCB system is overactive in obesity in peripheral organs and in brain regions involved in the control of energy balance such as the hypothalamus (for review: [18, 19]). However, this overactivation extends also to the hippocampus [20–22], where the eCB system controls synaptic plasticity and memory formation in healthy conditions [23–26]. The main eCBs expressed in the brain are anandamide (AEA) and 2-arachidonoyl glycerol (2-AG) and they represent primary ligands of the main cannabinoid receptor subtype 1 (CB_1_R) highly expressed on GABAergic interneurons and glutamatergic pyramidal neurons in the hippocampus [27, 28]. However, the exact mechanisms mediating the potential involvement of the eCB-CB_1_R system in HFD-induced memory deficits are currently unexplored.

A large body of evidence suggests strong interactions between the eCB system and the mechanistic Target of Rapamycin (mTOR) pathway in both lean and obese animals [29–34]. In particular, mTOR is a crucial molecular complex involved in multiple processes including nutrient sensing, synaptic plasticity and memory (for reviews: [35–37]). Thus, like the eCB system, mTOR appears to be at the crossway between obesity and cognition. For instance, whereas activation of mTOR pathway underlies amnestic effects of exogenous CB_1_R activation ([29, 30]), HFD alters mTOR signalling in several brain regions including the hippocampus [38] and blockade of the mTOR pathway prevents peripheral alterations in HFD-fed rodents (for review: [39]). However, the idea that the mTOR pathway, through a potential interaction with CB_1_R, might mediate memory deficits induced by HFD intake has never been explored.

Here we aimed to characterize the role of the eCB-CB_1_R system in memory deficits induced by obesogenic HFD consumption. We found that, after training, HFD enhances AEA levels and CB_1_R expression as well as CB_1_R-dependent neuronal activation, long-term potentiation and mTOR phosphorylation in the hippocampus. These events are causally linked, because the pharmacological blockade of eCB or mTOR signalling, as well as genetic deletion of CB_1_R in hippocampal glutamatergic neurons rescued HFD-induced long-term memory deficits. Therefore, these results demonstrate that diet-induced alterations of hippocampal eCB-CB_1_R system are responsible for aberrant hippocampal activity and plasticity leading to memory consolidation deficits.

## Materials and methods

### Animals, diet and housing

Male wild-type C57BL/6N mice (Janvier, France) or male *CB_1_R*-floxed mice (obtained, maintained and genotyped as described in Marsicano et al., 2003; Monory et al., 2006) aged 3 weeks at arrival were housed in groups of 8 individuals per cage in an air-conditioned room (22 ± 1°C) maintained under a 12 h light/dark cycle (lights on at 7:30 A.M., lights off at 7:30 P.M.). Food provided since their arrival was either standard laboratory chow, control diet (CD; A04 SAFE, Augy, France; offering 3.3 kcal/g consisting of 60% carbohydrate mostly from starch, and 3% fat), or high-fat high-sugar diet [HFD, D12451, Research Diets, New Brunswick, NJ; offering 4.7 kcal/g consisting of 24% fat (45% kcal), mostly saturated fat from lard, and 41% carbohydrate (35% kcal), with half coming from sucrose]. Mice had *ad libitum* access to water and food (either CD or HFD) during 12 weeks before starting the experiments. All the mice were regularly weighed from weaning until sacrifice and EchoMRI was used in a subset of mice (CD: n= 30; HFD: n= 29) to measure fat mass before sacrifice. All experiments were performed during the first part of the light phase (from 9am to 1pm). All animal care and experimental procedures were performed in accordance with standard ethical guidelines (European Directive 2010/63/UE). Experiments were approved by the local ethical committee (CEEA50) and the French Ministry of Agriculture and Forestry (authorization numbers APAFIS #15803, #30052 and #51264).

### Drugs

The CB_1_R antagonist Rimonabant (0.5 and 1 mg/kg; Cayman Chemical, Michigan, USA), and the mTOR inhibitor Temsirolimus (1 mg/kg, LC Laboratories, Woburn, Massachusetts, USA) were dissolved in a mixture of 4% ethanol, 4% Cremophor-EL and 92% of saline (NaCl 0.9%). A mixture of 4% ethanol, 4% Cremophor-EL and 92% of saline served as vehicle injection. All drugs were injected intraperitoneally (i.p.) in a volume of 10 ml/kg.

### Object recognition memory

Mice were handled once per day for 3 consecutive days in their home-cage room before starting the experiment. Object recognition memory (ORM) task was performed as previously described in mice [11, 40]. The experimental apparatus used for the object recognition task was an open-field box (40 cm x 40 cm) with 16 cm high walls made of white-painted wood, placed in the center of an experimental room (light intensity of 15 Lux). During training, mice were placed in the experimental apparatus, facing the wall, at the opposite end from the objects. Two identical objects (A1 and A2) were placed 8 cm from the walls and ∼13 cm apart. The objects were Lego^®^ brick towers, lab bottles, culture flasks or cylinders. The session duration was 10 min maximum in which mice were allowed to freely explore the objects until they reached a criterion of 20 seconds of total exploration of both objects, which reduces inter-individual variability as previously validated [40]. Animals that did not reach the 20-seconds criterion of total exploration in 10 minutes were excluded from the analysis. Time spent exploring each object was recorded. Object exploration, defined as nose and whiskers touching and/or pointed towards the object in a distance of less than 1–1.5 cm away, was manually quantified by a trained experimenter outside the experimental room blind to experimental groups. A video tracking system (Smart, Panlab, Barcelona, Spain) recorded the exact track of each mouse as well as total distance travelled (cm). After training, mice were removed from the apparatus and returned to their home-cage. To avoid the presence of olfactory trails, the objects and arena of open-field were thoroughly cleaned with 10% ethanol after each mouse.

Object recognition memory (ORM) was tested either 3 hours (short-term memory) or 24 hours (long-term memory) after the training trial. During the test trial, mice were placed in the same arena in which one copy of the familiar object (A3) and a new object (B) were placed in the same location as stimuli during the training trial. All combinations and locations of objects were used in a counterbalanced manner to reduce potential biases due to preference for particular location or object. As previously, mice were left to explore both objects until they reached a criterion of 20 seconds [40]. Time spent exploring each object was recorded and a similar cutoff of 10 minutes was established. Some animals received an i.p. injection of vehicle, CB_1_R antagonist (Rimonabant, 0.5 or 1 mg/kg) or mTOR inhibitor (Temsirolimus, 1 mg/kg), immediately after training and were tested 24 hours later.

### Brain c-Fos activation

Mice were given injections of a lethal dose of pentobarbital sodium (1 ml, i.p.) 90 minutes after ORM training (8 minutes) immediately followed by i.p. injection of vehicle or CB_1_R antagonists (1 mg/kg). They were then quickly perfused with 4% paraformaldehyde (PFA) in PBS. Brain were removed and stored at 4°C in 4% PFA for 24 h to allow post-fixation. The next day, they were submerged in 30% sucrose solution at 4°C for 48 h to allow cryoprotection. Finally, brains were stored at −80°C. For c-Fos immunoreactivity, coronal sections of 40 µm were incubated in PBS containing 3% bovine serum albumin (BSA) and 0.3% Triton 100X (blocking buffer) to block nonspecific binding sites and to facilitate antibody penetration. The sections were also saturated with 0.3% hydrogen peroxide for 30 min to eliminate endogenous peroxidase. Section were then incubated with the primary anti c-Fos antibody (anti c-Fos Rabbit polyclonal antibody diluted 1: 1000 in blocking buffer; Santa Cruz Biotechnology) for 24 h at 4°C and for 2 h with the biotinylated secondary antibody (goat anti-rabbit antibody, diluted 1:1000 in PBS; Vector Laboratories) at room temperature, followed by 1 h incubation in the avidin–biotin–peroxidase complex solution (Vectastain, diluted 1:1000 in PBS; Vector Laboratories). Between each treatment, sections were thoroughly rinsed with PBS. The peroxidase complex was visualized after incubation for 9 min in a mix containing diaminobenzidine, ammonium chloride, ammonium sulfate, sodium acetate, glucose, and glucose oxidase. Sections were incubated in sodium acetate to stop the enzymatic reaction, rinsed in PBS, mounted on gelatin-coated slides, dehydrated, and cover-slipped. Labeling was quantified bilaterally on three to four sections spaced 240 µm apart and chosen to cover the basolateral amygdale (BLA), the perirhinal cortex (PRH) and the different subfield of the dorsal hippocampus, CA1, CA3 and dentate gyrus (DG; 1.46-2.18 mm posterior to bregma according to Paxinos and Franklin, 2001). Each section was photographed (Nikon-ACT-1 software), and labeled cells were counted with ImageJ software on a surface representing 0.57 mm^2^. Results were expressed for mm^2^.

### In vivo hippocampal electrophysiology

Mice were anesthetized in a box containing 5% Isoflurane (VIRBAC, France) 15-20 minutes after being taken from home-cage or after ORM training (8 minutes) immediately followed for some animals by i.p. injection of vehicle or CB_1_R antagonist (1 mg/kg). They were then placed in a stereotaxic frame (Model 900, Kopf instruments, CA, USA) in which 1.0% to 1.5% of Isoflurane was continuously supplied via an anesthetic mask during the complete duration of the experiment. The body temperature was maintained at ∼36.5°C using a homeothermic system (model 50-7087-F, Harvard Apparatus, MA, USA) and the complete state of anesthesia was assured through a mild tail pinch. Before surgery, 100 µl of the local anesthetic Lurocaine^®^ (Vetoquinol, France) was injected in the scalp region. Surgical procedure started with a longitudinal incision of 1.5 cm in length aimed to expose Bregma and Lambda. After ensuring correct alignment of the head, two holes were drilled in the skull for electrode placement. Recordings of field excitatory postsynaptic potentials (fEPSPs) were made from the stratum radiatum in the CA1 region of the right hippocampal hemisphere in response to stimulation of the Schaffer collateral /commissural pathway. A glass recording electrode, inserted in the CA1 stratum radiatum, and one concentric bipolar electrode (Model CBARC50, ME, USA) in the CA3 region using the following coordinates: 1) CA1 stratum radiatum: 1.5 mm posterior, 1.0 mm lateral and 1.20 mm below Bregma according to Paxinos and Franklin, 2001; CA3: 2.2 mm posterior, 2.8 lateral and 1.3 mm below Bregma (20° insertion angle). The recording electrode (tip diameter = 1–2 μm, 4–6 MΩ) was filled with a 2% pontamine sky blue solution in 0.5M sodium acetate. At first the recording electrode was slowly lowered by hand until it reached the surface of the brain and then to the final depth using a hydraulic micropositioner (Model 2650, KOPF instruments, CA, USA). The stimulation electrode was placed in the correct area by hand using a standard manipulator. Both electrodes were adjusted to find the area with maximum response, i.e. when the appearance of a negative deflecting fEPSP with a latency of approximately 10 ms was observed. *In vivo* recordings of evoked fEPSPs were amplified 1000 times and filtered (low-pass at 1Hz and high-pass 3000Hz) by a DAGAN 2400A amplifier (DAGAN Corporation, MN, USA). fEPSPs were digitized and collected on-line using a laboratory interface and software (CED 1401, SPIKE 2; Cambridge Electronic Design, Cambridge, UK). Test pulses were generated through an Isolated Constant Current Stimulator (DS3, Digitimer, Hertfordshire, UK) triggered by the SPIKE 2 output sequencer via CED 1401 and collected every 2 seconds at a 10 kHz sampling frequency and then averaged every 180 seconds. Test pulse intensities were typically between 40-250 µA with a duration of 50 µseconds. Basal stimulation intensity was adjusted to 30-50% of the current intensity that evoked a maximum field response. All responses were expressed as percent from the average responses recorded during the 15 minutes before high-frequency stimulation. High-frequency stimulation was induced by applying 3 trains of 100 Hz (1 second each), separated by 20 seconds interval. fEPSP were then recorded for a period of 60 minutes. In the end of each experiment, the position of the electrodes was marked (recording area: iontophoretic infusion of the recording solution during 180s at −20µA; stimulation area: continuous current discharge over 20 seconds at +20µA) and histological verification was performed *ex vivo*.

### Hippocampal endocannabinoid quantification

Two cohorts of mice were used for the measurement of eCB levels in the hippocampal tissue at different time points after ORM training for both CD and HFD groups. Hippocampal tissues were rapidly isolated, weighed and frozen immediately (first cohort), 30 minutes and 60 minutes (second cohort) after ORM training (8 minutes). The training duration was similar for all mice in order to standardize environmental stimulations and to give sufficient time to investigate both objects for at least 20 seconds. Home-cage mice from CD and HFD groups were added to each cohort.

The extraction and quantification of the main eCBs, N-arachidonoyl ethanolamine (anandamide; AEA) and 2-arachidonoylglycerol (2-AG), from hippocampus were performed as described previously [41]. First, for each mouse, both hippocampi were homogenized and lipids extracted with chloroform /methanol /Tris-HCl 50 mM pH 7.5 (2: 1: 1, v/v) containing internal deuterated standards. Samples were then analyzed by liquid chromatography-tandem mass spectrometry (LC-MS/MS). Mass spectral analyses were performed on a TSQ Quantum triple quadrupole instrument (Thermo-Finnigan, San Jose, CA, USA) equipped with an atmospheric pressure chemical ionization source and operating in positive ion mode. A sensitive and specific LC-MS/MS method was developed and validated for AEA and 2-AG, the quantification of which was achieved by isotope dilution using a calibration curve. AEA and 2-AG levels after ORM training were expressed as % of their internal home-cage control as previously reported [42].

### Hippocampal CB_1_R expression

Home-cage mice or mice submitted to the ORM training (8 minutes) 60 minutes earlier were deeply anesthetized with isoflurane. Hippocampus was collected on ice in RNAse free conditions then frozen and stored at −80◦C. Total RNA was extracted using TRIzol Reagent™ (Invitrogen, France). Hippocampal tissue was homogenized with 1mL of TRIzol with a Tissue Lyser during 3 minutes at 30Hz (Qiagen) and chloroform was added to separate the organic and water phases. After 15 min centrifugation at 12000g and 4°C, the supernatant water phase containing total RNAs was recovered and the organic phase containing DNA and proteins was stored at −20°C. RNAs were precipitated by adding isopropanol and centrifuged, then the RNA pellet was washed with 75% ethanol. Finally, they were dissolved in sterilized water. RNA concentration purity and integrity were obtained with Nanodrop spectrophotometer (NanoDrop™ technologies, Fisher Scientific) and RNA 6000 Nano LabChip kit for 2100 Bioanalyzer (Agilent Technologies). Then, 1 µg of RNA were reverse transcribed into cDNAs using random primers and Transcriptase inverse SuperScript IV™ (Fisher scientific). Gene amplification was performed using a SYBR Green I Master kit (ref RR420L, TAKARABIO Europe, France) and appropriate primers for CB_1_R gene, *Cnr1* (Forward primer: CCTGGGAAGTGTCATCTTTGTCT; Reverse primer: GGTAACCCCACCCAGTTTGA). The real-time PCR was operated with the LightCycler 480 system (Roche Diagnostics) and β*-actin* served as housekeeping gene since its expression level was constant in all experimental conditions. mRNA relative quantification was obtained using the GenEx™ MultiD software, as previously described [15, 43].

### Stereotaxic surgery and viral injection

#### CB_1_R deletion

After 7 weeks of HFD consumption (i.e. 10 weeks-old mice), LoxP-flanked *Cnr1* transgenic mice (*CB_1_R*-floxed; [44]) were anesthetized by i.p. injection of a mixture of ketamine (100 mg/kg, Imalgene 500®, Merial, France) and Xylazine (10 mg/kg, Rompun, Bayer, France) and placed into a stereotaxic apparatus (David Kopf Instruments, France) with mouse adaptor and lateral ear bars. Specific hippocampal deletion of the *Cnr1* gene (encoding CB_1_R protein) were obtained by bilateral injection of adeno-associated viruses expressing the Cre recombinase under a general promoter (AAV8/2-CAG-EGFP_Cre-WPRE-SV40p(A), titre 2.1 x 10E12 vg/ml and its control AAV8/2-CAG-EGFP-WPRE-SV40p(A), titre 2.9 x 10E12 vg/ml; ETH VVF, Zurich), a specific promoter for glutamatergic projecting neurons (AAV1/2-mCaMKIIα-EGFP_2A_iCre-WPRE-SV40p(A), titre 4.6 x 10E12 vg/ml and its control AAV1/2-mCaMKIIα-mCherry-WPRE-hGHp(A), titre 4.8 x 10E12 vg/ml; ETH VVF, Zurich) or a specific promoter for GABAergic interneurons (AAV-9/2-mDlx-HBB-chI-iCre-WPRE-bGHp(A), titre 5.3 x 10E11 vg/ml; ETH VVF, Zurich) into the hippocampus (2 injections of 1 µl per side). AAV vectors were injected with the help of a microsyringe attached to a pump (UMP3-1, World Precision Instruments) at the following coordinates: −2 mm antero-posterior, ±1.5 mm medio-lateral from Bregma, −2 mm (first injection) and −1.5 mm (second injection) dorso-ventral from the skull surface, according to Paxinos and Franklin, 2001. Animals remained on HFD after surgery until sacrifice. ORM training with test at 24 hours was performed 5 weeks after injections in order to get an optimal *Cnr1* deletion in the hippocampus.

#### Viral expression

In order to verify hippocampal GFP or mCherry expression, mice were given injections of a lethal dose of pentobarbital sodium (1 ml, i.p.) and quickly perfused with 4% paraformaldehyde (PFA) in PBS as described previously. Coronal sections of 40 µm were rinsed four times in Tris-NaCl 0.05M (30 min each rinse) to eliminated cryoprotector solution. The sections were then mounted on slides with mounting medium to preserve fluorescence-stained tissue sections (Calbiochem, Billerica, MA, USA) and coverslipped. The sections were visualized with fluorescent microscope Nikon Eclipse E400 at 10x magnification and were photographed using NIS-Element AR 3.2 software.

#### Quantification of CB_1_R expression

In order to evaluate CB_1_R expression and to quantify the decrease in animals injected with Cre recombinase, another set of floating slices were incubated with a guinea pig anti-CB_1_R antibody (1:1000, Cell Signaling Technology) for 24 h at 4°C. Slices were then incubated for 2 h at room temperature with a goat anti-guinea pig antibody (A594; 1:2000, Vector Laboratories). Slices were then mounted on gelatin-coated slides and coverslipped. All slices were photographed with Nanozoomer slide scanner Hamamatsu NANOZOOMER 2.0HT (Bordeaux Imaging Center, University of Bordeaux, France). QuPath program (QuPath v.0.3.0 40) was used for quantification of CB_1_R density (number of CB_1_R/µm²). Labeling was quantified in the dorsal hippocampus and the basolateral amygdala as a control brain area.

### Protein extraction and western blot

Two independent experiments were performed using different protein extraction and western blot protocols (see below). The two experiments leading to similar results, the data were thus pooled. For a first experiment, total proteins from hippocampi were extracted from the TRIzol fraction previously recovered from the RNA extraction step. After several steps of precipitation and washing the protein, pellet was solubilized by sonication in a 1:1 solution of 1% SDS and 8 M urea in Tris-HCl 1M, pH 8.0 and stored at −80°C. Protein concentration was determined using the MicroBC Assay protein quantification kit. Western blots were performed to quantify several proteins implicated in the downstream activation pathways of CB_1_R: mTOR, phospho-mTOR, MAPK/ERK1/2 and phospho-MAPK/ERK1/2. For the Western blot analysis 25 µg of proteins were loaded on a polyacrylamide gel 10% and then separated by electrophoresis (for 1h30). After that, proteins were transferred (for 1h30) on a nitrocellulose membrane (0.2 µm, Amersham Protran). Unspecific sites were saturated by a solution of TBS/Tween20 0.1%-milk 5%. Membranes were then incubated with a primary antibody solution diluted in TBS-Tween/Na azide/BSA: anti-beta-actin, as a reference (1/5000, Merck), anti-mTOR (1/1000 Cell Signaling), anti-phospho-mTOR (1/1000, Cell Signaling), anti-p44/42 MAPK/ERK1/2 (1/1000, Cell Signaling) and anti-phospho-p44/42 MAPK/ERK1/2 (Thr202/Tyr204; 1/1000, Cell Signaling). After one night of incubation at 4°C, membranes were washed with a TBS-Tween solution and incubated with the secondary antibody anti-rabbit coupled with HRP (1/5000) for 1h at room temperature. Finally, the immunostaining was revealed by incubating the membrane with Luminata Crescendo (Millipore). Band signal was quantified using the Chemidoc MP (BioRad) and analyzed using the ImageLab Chemidoc software (BioRad). The signal for each protein was normalized to the β-actin signal intensity for each sample.

In a second experiment, additional samples were processed using fluorescent instead of chemiluminescent detection. To extract proteins, hippocampi were homogenized using Tissue lyser (Qiagen) for 15s at 30Hz in 50mM Tris, 2% SDS, 0.5M Urea and supplied with anti-phosphatase cocktail then homogenates were sonicated using Bioruptor PLUS (Diagenode) for 6 cycles of 30s ON / 30s OFF. Homogenates were incubated on ice for 30 minutes and centrifuged for 5 minutes at 3600rpm. Proteins concentration was then measured using Micro BC Assay Uptima (Interchim). For the Western blot analysis 20 µg of denaturated proteins were loaded on a polyacrylamide Mini-PROTEAN Precast gel 4-20% (Biorad). Electrophoresis and transfer on nitrocellulose membranes were performed as described above. Unspecific sites were saturated by LICOR Intercept TBS blocking buffer (LICOR). Membranes were then incubated with a primary antibody solution diluted in TBS-1% milk: mouse anti-alpha-Tubulin, as a reference (1/5000, Merck), rabbit anti-phospho-mTOR (1/1000, Cell Signaling) and rabbit anti-phospho-p44/42 MAPK/ERK1/2, (Thr202/Tyr204; 1/1000, Cell Signaling). After one night of incubation at 4°C, membranes were washed with a TBS solution and incubated with a mix of secondary antibodies, anti-mouse coupled with IRDye 700 and anti-rabbit coupled with IRDye 800 (1/10000) for 1h at room temperature. Alpha-tubulin and phosphorylated protein forms immunostaining fluorescence was recorded onto Odissey apparatus. Band signal was quantified and analyzed using the Image Studio software. Membranes were incubated 10 minutes into Re-blot Plus Strong solution (Millipore) to discard antibodies against alpha-tubulin and phosphorylated protein forms then membranes were incubated overnight at 4°C with rabbit anti-mTOR (1/1000, Cell Signaling), rabbit anti-p44/42 MAPK/ERK1/2 (1/1000, Cell Signaling), membranes were washed with a TBS solution and incubated with a secondary antibody anti-rabbit coupled with IRDye 800 (1/10000) for 1h at room temperature. Total protein forms immunostaining fluorescence was recorded onto Odissey apparatus. Band signal was quantified and analyzed using the Image Studio software. The signal intensity for the phosphorylated forms was normalised to the signal intensity of protein total forms.

### Statistics

Data are expressed as the mean ± SEM, and analyses were conducted using Prism software (GraphPad) with a threshold for considering statistically significant difference being set up at *p* ≤ 0.05. Comparison between groups were analysed using unpaired Student’s t-tests or one-two-or three-way ANOVA (with repeated measures when appropriate). When the interaction was significant, it was further investigated using *post-hoc* Fisher’s PLSD tests. Moreover, for each group, ORM performance was compared against reference level (50% exploration ratio for memory tasks or 100% baseline for LTP) using one-sample *t*-test.

## Results

### CB_1_R blockade rescues HFD-induced long-term memory deficits

We previously reported morphometric and metabolic changes induced by peri-adolescent HFD consumption in mice [11, 12, 15]. Here, we confirmed those results, showing that HFD consumption for 12 weeks from weaning to adulthood enhanced body weight (CD: 28.9 ± 0.3 g; HFD: 32.7 ± 0.5 g; unpaired t-test: *t*_(57)_ = 6.28, *P* < 0.0001) and fat mass (CD: 3.5 ± 0.2 g; HFD: 8.3 ± 0.6 g; *t*_(57)_ = 8.30, *P* < 0.0001).

Similar to what we recently found in rats [14], we observed that peri-adolescent HFD impaired long-term, but not short-term, ORM in mice assessed 24 hours and 3 hours after training, respectively (Figure 1A-C, Supplementary Table 1). Similar results were obtained using an L-maze instead of an open-field to assess short- and long-term ORM (Supplementary Figure 1 and Table 1). These findings indicate that peri-adolescent HFD consumption induces metabolic alterations and a specific impairment of ORM consolidation. As higher eCB activity was reported in obese subjects [18–22], we then evaluated the effect of blocking CB_1_R activation on long-term ORM. Systemic administrations of the CB1R antagonist Rimonabant immediately after training altered ORM performance in a dose dependent manner. While a low dose (0.5 mg/kg) did not alter performance in both CD and HFD-fed mice, strikingly, a higher dose (1 mg/kg) rescued HFD-induced ORM deficit and impaired ORM in CD-fed mice (Figure 1D, Supplementary Table 1).

**Figure 1.**
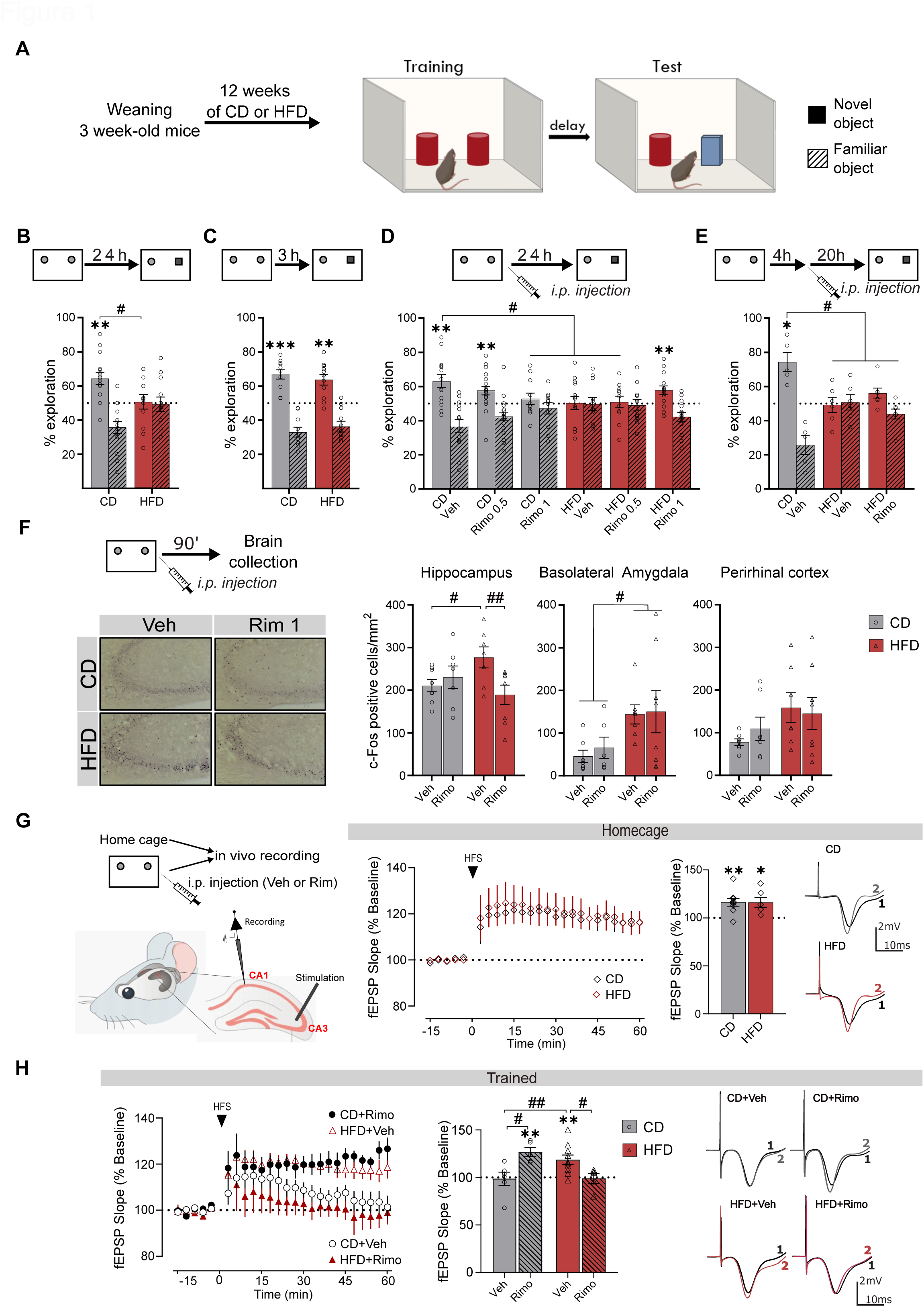
HFD exposure induces long-term memory deficits, training-induced higher hippocampal activity and aberrant hippocampal synaptic plasticity that are dependent on CB_1_R activation. (A) Schematic representation of object recognition memory task. (B, C) HFD exposure impairs long-term (B) but not short-term (C) object recognition memory. (D, E) HFD-induced impairment of long-term object recognition memory is rescued by systemic injection of 1 mg/kg, but not 0.5 mg/kg, CB_1_R antagonist (Rimonabant) immediately after training (D) but not 4 hours later (E). (F) Systemic injection of CB_1_R antagonist (Rimonabant 1 mg/kg) reverses HFD-induced higher Fos activation after training in the hippocampus but not in the amygdala or the perirhinal cortex. (G) HFD consumption does not affect *in vivo* long-term potentiation in CA3-CA1 pathway at baseline. (H) HFD-induced aberrant long-term potentiation in CA3-CA1 pathway after training is normalized by systemic injection of CB_1_R antagonist (Rimonabant 1 mg/kg). * *P* < 0.05, ** *P* < 0.01, *** *P* < 0.001: one-sample t-test (B-E: against 50%; G-H against 100%); # *P* < 0.05, ## *P* < 0.01: unpaired t-test (B); one-way or two-way ANOVA (D-F), two or three-way repeated measures ANOVA (G-H) followed by *post-hoc* test. For statistical details, see Supplementary Table 1.

Interestingly, delayed injection of 1 mg/kg Rimonabant 4 hours after training did not alleviate ORM impairment in HFD-fed mice (Figure 1E, Supplementary Table 1), indicating that over-activation of CB_1_R during a short time window after training is responsible for ORM consolidation deficits in HFD-fed mice.

### CB_1_R blockade normalizes HFD-induced hippocampal over-activation after training

To identify which brain structure(s) could be involved in the beneficial effects of CB1R blockade in HFD-fed animals, we assessed the expression levels of the marker of neuronal activity c-Fos 90 minutes after training. Based on their involvement in long-term ORM, we focused on the dorsal hippocampus (DG, CA1 and CA3), the perirhinal cortex and the basolateral amygdala (for reviews: [45, 46]). In home-cage animals, c-Fos expression was low in all these structures and not different between CD and HFD groups (data not shown; see [43]). As previously reported [11, 43], training induced a higher c-Fos expression in HFD-fed mice than CD-fed mice in all 3 structures. Systemic injections of the CB_1_R antagonist (at 1 mg/kg) did not alter this expression in the perirhinal cortex and the basolateral amygdala (Figure 1F, Supplementary Table 1). Interestingly, however, pharmacological CB1R blockade normalized c-Fos levels in the hippocampus of HFD-fed mice (Figure 1F, Supplementary Table 1), particularly in the CA3 area (Supplementary Figure 2 and Table 1). These results indicate that peri-adolescent HFD leads to training-induced c-Fos over-expression in several forebrain areas, but this effect is specifically under the control of CB_1_R activation in the hippocampus.

#### CB_1_R blockade normalizes HFD-induced aberrant hippocampal synaptic plasticity

Synaptic plasticity and in particular long-term potentiation (LTP) has been proposed as a cellular mechanism of long-term memory in the hippocampus (for review: [47]). We thus wondered whether HFD consumption since weaning altered *in vivo* hippocampal LTP through CB_1_R activation. In home cage animals, high-frequency stimulation induced stable and similar LTP of field excitatory post-synaptic potential (fEPSP) slopes in both CD and HFD groups one hour after stimulation (Figure 1G, Supplementary Table 1). These results indicate that adolescent HFD consumption does not affect CA3-CA1 LTP in home-cage mice.

Because LTP can be affected by ORM training [26, 48], we then investigated the impact of CB_1_R blockade on post-training LTP in CD and HFD-fed mice. Similar to the behavioral results described above, training and CB_1_R blockade affected CA1 LTP in opposite directions, depending on the diet. Thus, after training, LTP was absent in the CD-vehicle group, whereas high-frequency stimulation induced normal LTP in the HFD-vehicle group, indicating that diet reverts the effects of training on this form of hippocampal plasticity (Figure 1H, Supplementary Table 1). In CD-fed mice, the post-training administration of the CB_1_R antagonist abolished the inhibitory effect of training on LTP (Figure 1H, Supplementary Table 1). Conversely, Rimonabant restored the normal inhibitory effect of training on this form of *in vivo* synaptic plasticity in trained HFD-fed animals (Figure 1H, Supplementary Table 1). These results suggest that, in normal conditions, training suppresses hippocampal plasticity induced by high-frequency stimulation of the CA3-CA1 pathway. Peri-adolescent HFD fully inverts these phenomena, possibly underlying the cognitive deficit. Strikingly, CB_1_R blockade restores CD-like plasticity responses in trained HFD-fed animals.

#### HFD intake upregulates hippocampal eCB system after ORM training

We then investigated whether HFD intake affects hippocampal eCB levels in home cage and ORM trained mice (Figure 2A and B). In home-cage animals, hippocampal levels of eCBs did not differ between groups (Figure 2A, Supplementary Table 1). In ORM trained mice, the hippocampal AEA content greatly increased in both groups 30 min and 60 min after training, with a higher increase in HFD group as compared to CD group 30 min after training (Figure 2B left, Supplementary Table 1). Relative to the home cage group, hippocampal 2-AG content did not vary after ORM (Figure 2B right, Supplementary Table 1). Regarding CB_1_R expression, we found that ORM increased hippocampal CB_1_R mRNA expression in HFD-fed mice but not in CD animals (Figure 2C, Supplementary Table 1). A similar, though non-significant, trend was observed with CB_1_R protein (Supplementary Figure 3A and Table 1).

**Figure 2.**
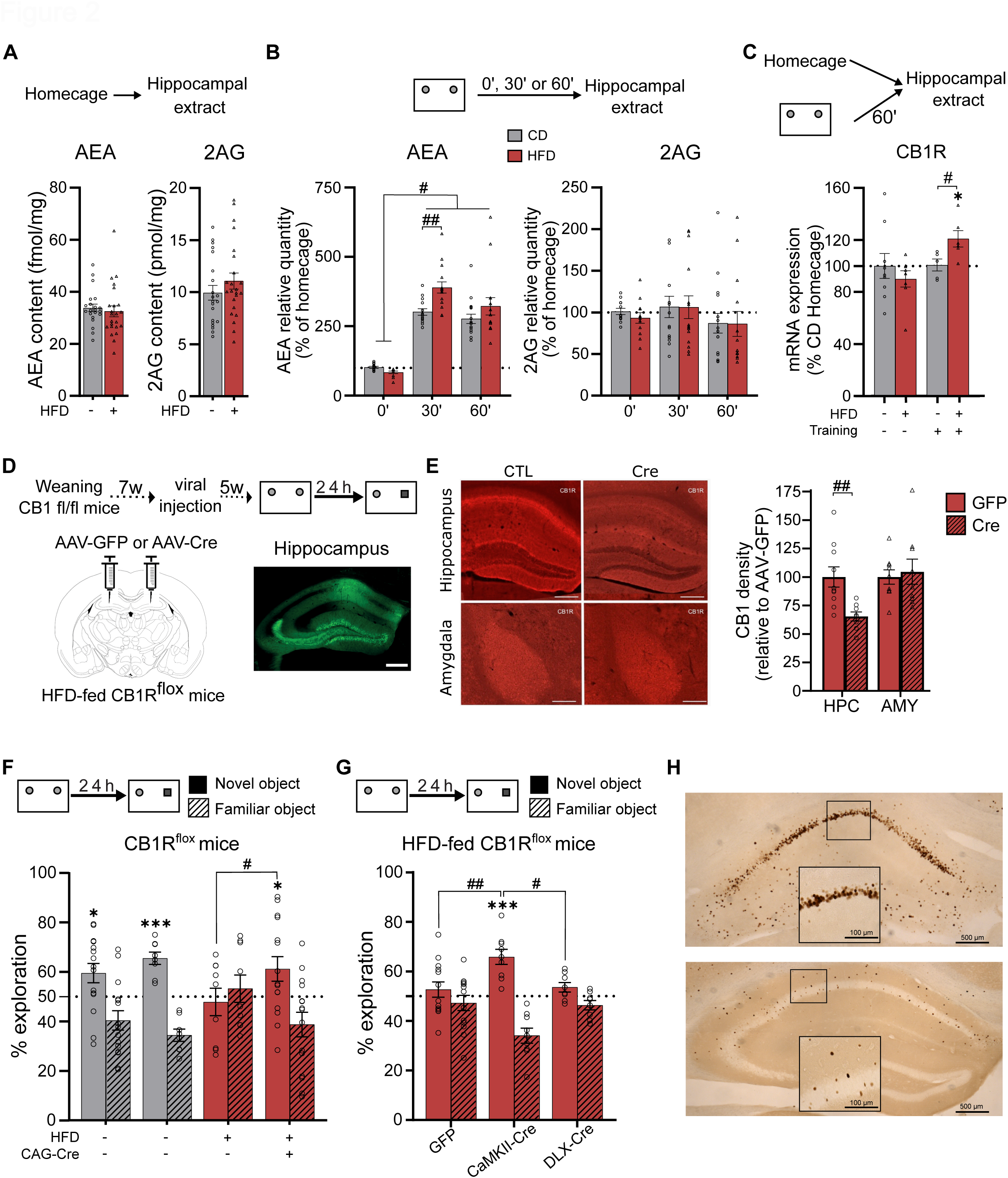
HFD exposure enhances training-induced anandamide levels and CB_1_R expression and hippocampal CB_1_R deletion rescues HFD-induced memory deficits. (A, B) HFD exposure does not affect anandamide (AEA) or 2-arachidonoylglycerol (2AG) hippocampal levels at baseline (A) but specifically enhances training-induced AEA hippocampal levels. Training does not change 2AG hippocampal levels in both groups. (C) HFD exposure enhances hippocampal CB_1_R mRNA expression after training but not at baseline. (D) Schematic representation of the protocol used to evaluate the effect of hippocampal CB_1_R deletion on object recognition memory task. (E, F) Viral expression of Cre-recombinase in the hippocampus of CB_1_R-floxed mice decreases by ∼35% CB_1_R expression specifically in the hippocampus, without changing CB_1_R expression in the amygdala (E), and rescues HFD-induced impairment of long-term object recognition memory (F). (G) Viral expression of Cre-recombinase driven by CaMKII promoter (targeting pyramidal glutamatergic neurons), but not Dlx promoter (targeting interneurons), in the hippocampus of CB_1_R-floxed mice rescues HFD-induced object recognition memory deficit. (H) Representative images of Cre immunostaining in the dorsal hippocampus of CB_1_R-floxed mice injected with either CaMKII-Cre (up) or Dlx-Cre (down) virus. Scale bar is set at 100 µm or 500 µm. * *P* < 0.05, *** *P* < 0.001: one-sample t-test (C: against 100%; F-G: against 50%); # *P* < 0.05, ## *P* < 0.01: unpaired t-test (C, E); one-way or two-way ANOVA (B, F-G), followed by *post-hoc* test. For statistical details, see Supplementary Table 1.

#### Deletion of CB_1_R in hippocampal glutamatergic neurons rescues long-term ORM in HFD-fed mice

Our findings support an overactivity of the hippocampal eCB system in HFD animals. We therefore evaluated whether the deletion of hippocampal CB_1_R could improve HFD-induced ORM impairment. We virally expressed the Cre recombinase (AAV-CAG-Cre) – or GFP as a control - in the hippocampus of HFD-fed *CB_1_R*-floxed mice (Figure 2D; [44]). This approach induced around 35% reduction of CB_1_R expression specifically in dorsal hippocampus (Figure 2E, Supplementary Table 1). Next, we evaluated the effect of this hippocampal *CB_1_R* downregulation on ORM performance. Whereas it did not affect ORM performance in CD-fed mice, it completely rescued ORM deficits in HFD-fed mice (Figure 2F, Supplementary Table 1). This indicates that the expression of hippocampal CB_1_R is required for HFD-induced ORM deficit in mice.

In the hippocampus, neuronal CB_1_R are abundantly expressed on GABAergic interneurons and glutamatergic pyramidal neurons [49, 50]. In order to evaluate which of these cell populations mediates CB_1_R-dependent memory deficits in HFD-fed mice, we virally expressed the Cre recombinase in dorsal hippocampal glutamatergic (using the CaMKII promoter, AAV-CaMKII-Cre [51]) or GABAergic neurons (using the Dlx promoter, AAV-Dlx-Cre [52]) of *CB_1_R*-floxed mice. Deletion of hippocampal CB_1_R on glutamatergic neurons rescued HFD-induced ORM impairment whereas lack of hippocampal CB_1_R from GABAergic neurons had no effect (Figure 2G-H and Table 1), despite similar decrease of CB_1_R expression in the hippocampus with both viruses (Supplementary Figure 3B and Table 1).

#### A CB_1_R–mTOR link mediates HFD-induced ORM deficit

Our data indicate that HFD impairs ORM consolidation by enhancing CB_1_R signaling on hippocampal glutamatergic neurons. Next, we wondered which downstream signaling pathways could be differentially recruited in the hippocampus of CD and HFD-fed mice. Based on their involvement in CB_1_R signaling, body energy metabolism and memory performance, we focused on the mTOR and MAPK/ERK pathways. Western blot analysis revealed that mTOR phosphorylation (phospho-mTOR) was not altered by HFD in untrained animals (Figure 3A left, Supplementary Figure 4A and Table 1). However, HFD significantly increased phospho-mTOR in ORM trained animals (Figure 3A left, Supplementary Figure 4A and Table 1). Conversely, MAPK/ERK phosphorylation (phospho-ERK) was unchanged in all conditions (Figure 3A right, Supplementary Figure 4A and Table 1). Next, we assessed whether CB_1_R blockade could normalize mTOR activation. Western immunoblotting revealed that the HFD-induced increase of phospho-mTOR in trained mice was abolished by a treatment with the CB_1_R antagonist Rimonabant (Figure 3B, Supplementary Figure 4B and Table 1).

**Figure 3.**
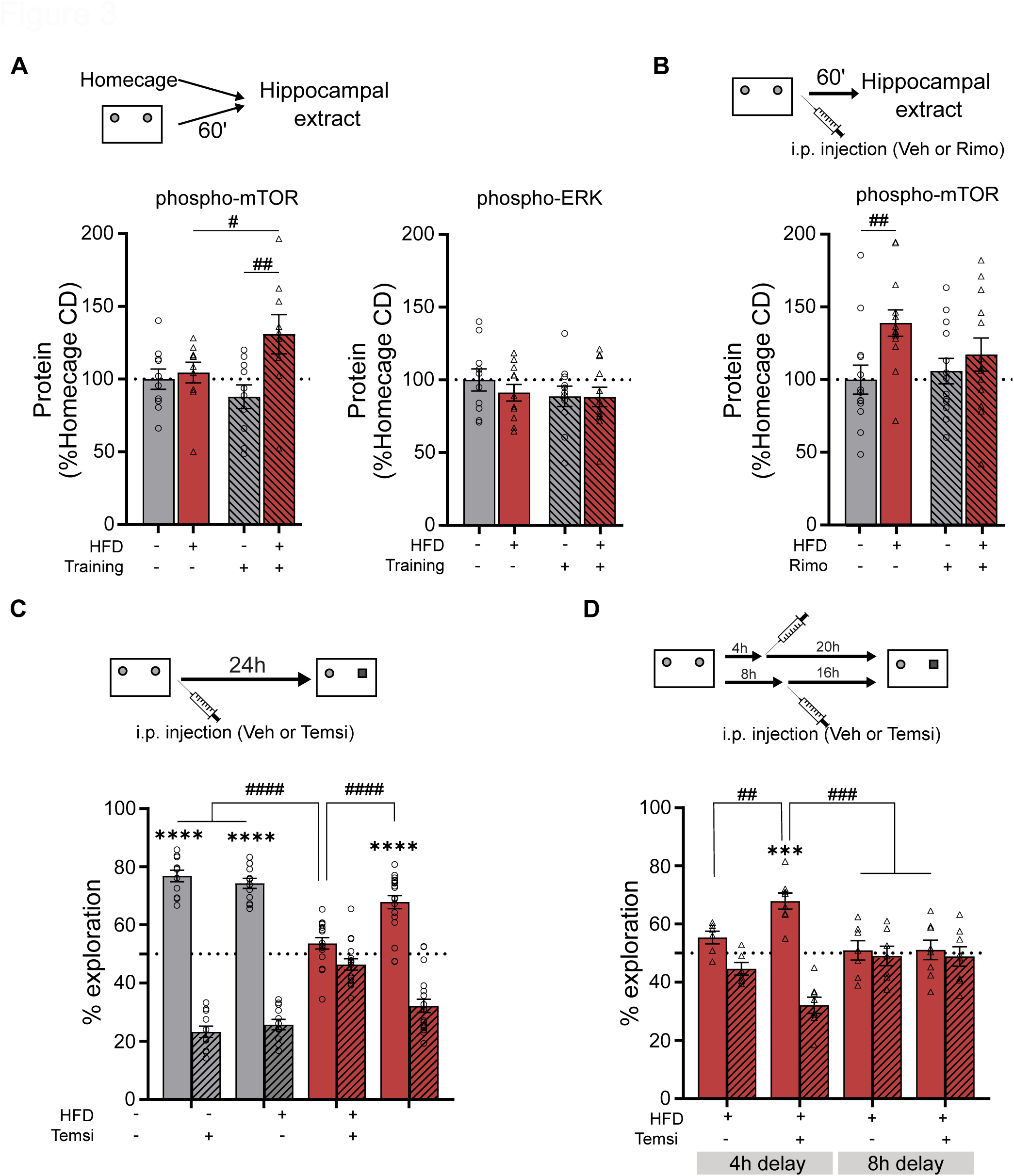
HFD exposure enhances training-induced mTOR phosphorylation and inhibiting mTOR rescues HFD-induced long-term memory deficits. (A) HFD exposure enhances hippocampal phosphorylation of mTOR after training, but not at baseline (left), without affecting hippocampal phosphorylation of ERK (right). (B) HFD-induced enhancement of hippocampal mTOR phosphorylation after training is blocked by systemic injection of CB_1_R antagonist (Rimonabant 1 mg/kg). (C-D) HFD-induced impairment of long-term object recognition memory is rescued by systemic injection of mTOR inhibitor (Temsirolimus 1 mg/kg) immediately after training (C) or 4 hours later, but not 8 hours later (D). *** *P* < 0.001, **** *P* < 0.0001: one-sample t-test (C-D: against 50%); # *P* < 0.05, ## *P* < 0.01, ### *P* < 0.001: two-way ANOVA followed by *post-hoc* test (A, B, C, D). For statistical details, see Supplementary Table 1.

Finally, in order to establish causal relationships between mTOR signaling and ORM impairment, we tested whether pharmacological inhibition of mTOR could improve ORM performance in HFD-fed mice. Notably, immediate post-training systemic injections of the mTOR inhibitor Temsirolimus (1mg/kg) fully rescued HFD-induced ORM deficits (Figure 3C, Supplementary Table 1). A similar beneficial effect was obtained with the injection of the mTOR inhibitor 4 hours, but not 8 hours, after training (Figure 3D; Supplementary Table 1). Thus, these results indicate that, in HFD-fed mice, training increases mTOR phosphorylation in a CB_1_R-dependent fashion, leading to impairment of ORM consolidation.

## DISCUSSION

This study reveals that diet-induced cognitive impairments are mediated by activation of the hippocampal eCB-CB_1_R system, through the modulation of neuronal activity, synaptic plasticity and mTOR signalling in mice.

### HFD consumption impairs ORM consolidation and induces aberrant hippocampal activity and plasticity through CB_1_R activation

The consumption of HFD during 12 weeks since weaning impairs long-term but not short-term ORM in mice, as we recently demonstrated in rats [14]. Together with previous results on different forms of memory [13], the present results suggest that HFD intake during adolescence specifically affects cellular mechanisms underlying consolidation processes. Based on the evidence that the eCB system is overactive in obesity (for review: [18, 19]), we showed that systemic CB_1_R blockade immediately, but not 4 hours, after training prevented HFD-induced long-term ORM impairment in a dose-dependent manner.

ORM training also induces higher c-Fos expression in the hippocampus, the perirhinal cortex and the basolateral amygdala in HFD-fed mice, as compared to CD-fed mice. HFD-induced increase of neuronal activity has been functionally linked to impaired memory [11, 14, 15]. Systemic blockade of CB_1_R immediately after training normalized c-Fos expression in the dorsal hippocampus, particularly in CA3, but not in the perirhinal cortex and the basolateral amygdala, indicating a specific CB_1_R modulation of hippocampal activity in HFD-fed mice. This indicates that the impact of HFD on the activity of CB_1_R is not generalized, but it is linked to specific brain systems and specific neuronal population (i.e. hippocampal glutamatergic neurons; see below). In this sense, it is important to note that HFD-fed female mice respond to training with different patterns of c-Fos activation than males [15]. Since the present experiments were performed in male mice, it will be important in future experiments to investigate potential sex-differences in the mechanisms here described.

Our data show that HFD affects *in vivo* synaptic plasticity in the dorsal hippocampus of anesthetized mice. Whereas the HFD did not alter CA3-CA1 LTP under basal conditions, ORM training impeded LTP in CD-fed mice suggesting that, in our conditions, training triggered hippocampal plasticity able to interfere with subsequent LTP. This interference could be due to either an occlusion of LTP or a depotentiation. Indeed, it was demonstrated that exposure to novel objects is followed by an enhancement of synaptic efficacy in CA3-CA1 synapses [26, 48]. This could indeed subsequently occlude CA3-CA1 LTP. It was also reported that ORM training also triggers another type of synaptic plasticity, e.g. hippocampal CA1 long-term depression [53, 54] which subsequently could interfere with CA1 LTP through depotentiation [55]. The detailed mechanisms involved in these complex synaptic interactions remain to be characterised, but our data show that these processes are fully reverted by the exposure to HFD for 12 weeks starting at weaning. One of the most striking results of this paper is that CB_1_R blockade reverts training-induced plasticity of HFD-fed mice to one that is undistinguishable from CD-fed animals, thereby establishing a very strong link with the behavioral results.

Interestingly, the same dose of CB1R antagonist (1 mg/kg) that normalized *in vivo* synaptic plasticity in the dorsal hippocampus and improved ORM in the HFD group (1 mg/kg), altered *in vivo* synaptic plasticity in the dorsal hippocampus and impaired ORM in the CD-fed group. This suggests that the eCB-CB_1_R system modulates hippocampal plasticity and memory in a state-dependent manner. In this case, the alterations caused by HFD appear to fully invert the impact of this signaling on plasticity and memory consolidation. This is in agreement with the complex and multifaceted effects of (endo)cannabinoid signaling in hippocampal synaptic plasticity and episodic-like recognition memory [26, 29, 42, 56–58].

### Hippocampal eCB-CB_1_R system mediates HFD-induced ORM deficits through mTOR pathway

ORM training clearly increases AEA, but not 2-AG, hippocampal levels independently of diet, suggesting that hippocampal AEA plays a critical role in modulating ORM. Selective increase of AEA levels has also been reported in the hippocampus after aversive learning in lean animals, suggesting that this eCB could be particularly important to regulate memory formation [59]. Interestingly, HFD-fed mice display higher AEA hippocampal levels than CD-fed mice 30 minutes after training, corroborating previous results [22, 60]. This finding is consistent with a previous study showing that enhancing the concentration of hippocampal AEA, but not 2-AG, impairs ORM in lean mice. This suggests that supra-physiological AEA levels after training are associated with ORM impairment, as in the case of HFD-fed mice. In addition to AEA, ORM training enhances CB_1_R expression and CB_1_R-dependent mTOR phosphorylation in the hippocampus of HFD-fed mice only. Interestingly, previous studies demonstrated that exogenous CB_1_R activation triggers amnesia through a similar activation of the mTOR pathway [29, 30], and our genetic and pharmacological data clearly show that both hippocampal CB_1_R and mTOR signalling are necessary for the impairment of memory consolidation induced by HFD.

Our data indicate that CB_1_R expressed in hippocampal glutamatergic, but not GABAergic, neurons are responsible for the HFD-induced long-term ORM deficit. Similarly, pharmacological CB_1_R activation induced amnesia of object location, another hippocampal-dependent memory task, through CB1R expressed on glutamatergic, but not GABAergic, neurons [62]. However, another study using long-term ORM reported that CB1R specifically expressed on GABAergic neurons mediates memory deficits induced by exogenous CB_1_R activation [29]. The discrepancy between this and our study could be due to the use of differential metabolic states (lean or obese) or differential CB1R cerebral manipulation (developmental and global through genetic deletion vs focal at adulthood through combination of genetic and viral approaches). This result adds to the complex and multifaceted regulation of recognition memory by CB_1_R. For instance, certain populations of CB_1_R are required for proper physiological ORM consolidation, but they are also responsible of ORM impairment when overactivated [58].

## Conclusion

This study reveals a novel mechanism underlying the impact of HFD on cognition. HFD-induced increase of hippocampal eCBs, activation of CB_1_R in glutamatergic neurons and stimulation of mTOR leading to maladaptive neuronal activation and synaptic plasticity represent a reasonable molecular and cellular cascade able to explain how metabolic alterations can be translated into cognitive ones. Teenager obesity reached the definition of epidemy worldwide, and it dramatically impacts the cognitive development of young people. Therefore, by revealing the mechanisms linking obesogenic diet and memory impairments, these data will likely bear important consequences on human and public health.

## Acknowledgements

This work was supported by INRAE (to G.F.), INSERM (to G.M.), Thai Ministry of Higher Education, Science, Research and Innovation (to Y.J.), Fondation pour la Recherche Medicale (FDT20160435664, to J.F.O.C), and French State/Agence Nationale de la Recherche (OBETEEN ANR-15-CE17-0013 to G.F.; ORUPS ANR-16-CE37-0010 to G.M. and G.F; HIPPOBESE ANR-23-CE14-0004 to G.F. and G.M.). We thank Helen McMillan, Domitille Rajot, Marianela Santoyo-Zedillo and Fabien Naneix for technical assistance. We also thank all the personnel of the animal facility of the NutriNeuro lab for mouse care and the CIRCE (Behavioural Engineering Centre) facility of Bordeaux Neurocampus.

---

**Supplementary Figure 1**

(A) Schematic representation of object recognition memory task using L-maze apparatus. (B-D C) HFD exposure impairs long-term (B) but not short-term (C) object recognition memory without affecting the total time of objects exploration during test (D).

* *P* < 0.05, ** *P* < 0.01, *** *P* < 0.001: one-sample t-test (against 50%); ## *P* < 0.01: unpaired t-test. For statistical details, see Supplementary Table 1.

**Supplementary Figure 2**

(A) Schematic representation of the protocol used to assess the effect of systemic injection of CB_1_R antagonist (Rimonabant 1 mg/kg) on training-induced Fos activation. (B-D) Rimonabant does not affect Fos expression in the dentate gyrus (DG; B) and the CA1 (C) but reversed HFD-induced higher Fos activation after training in the CA3 of the hippocampus (D).

\# *P* < 0.05: two-way ANOVA followed by *post-hoc* test. For statistical details, see Supplementary Table 1.

**Supplementary Figure 3**

(A) HFD exposure effect on hippocampal CB_1_R protein expression at baseline and after training assessed using Western-blot. (B) Both CaMKII-Cre and Dlx-Cre viruses decrease hippocampal CB_1_R expression by ∼15% (H).

**Supplementary Figure 4**

(A) Representative immunoblot of phospho-mTOR and phospho-ERK levels in homecage or trained mice fed control diet (CD) or HFD. (B) Representative immunoblot of phospho-mTOR and phospho-ERK levels in CD and HFD-fed mice injected immediately after training with vehicle or CB_1_R antagonist (Rimonabant 1 mg/kg, i.p.).

