## Supplementary figures and images for "Obesogenic diet impairs memory consolidation via the hippocampal endocannabinoid system"

### Supplemental Figure 1

#### Supplementary Figure 1

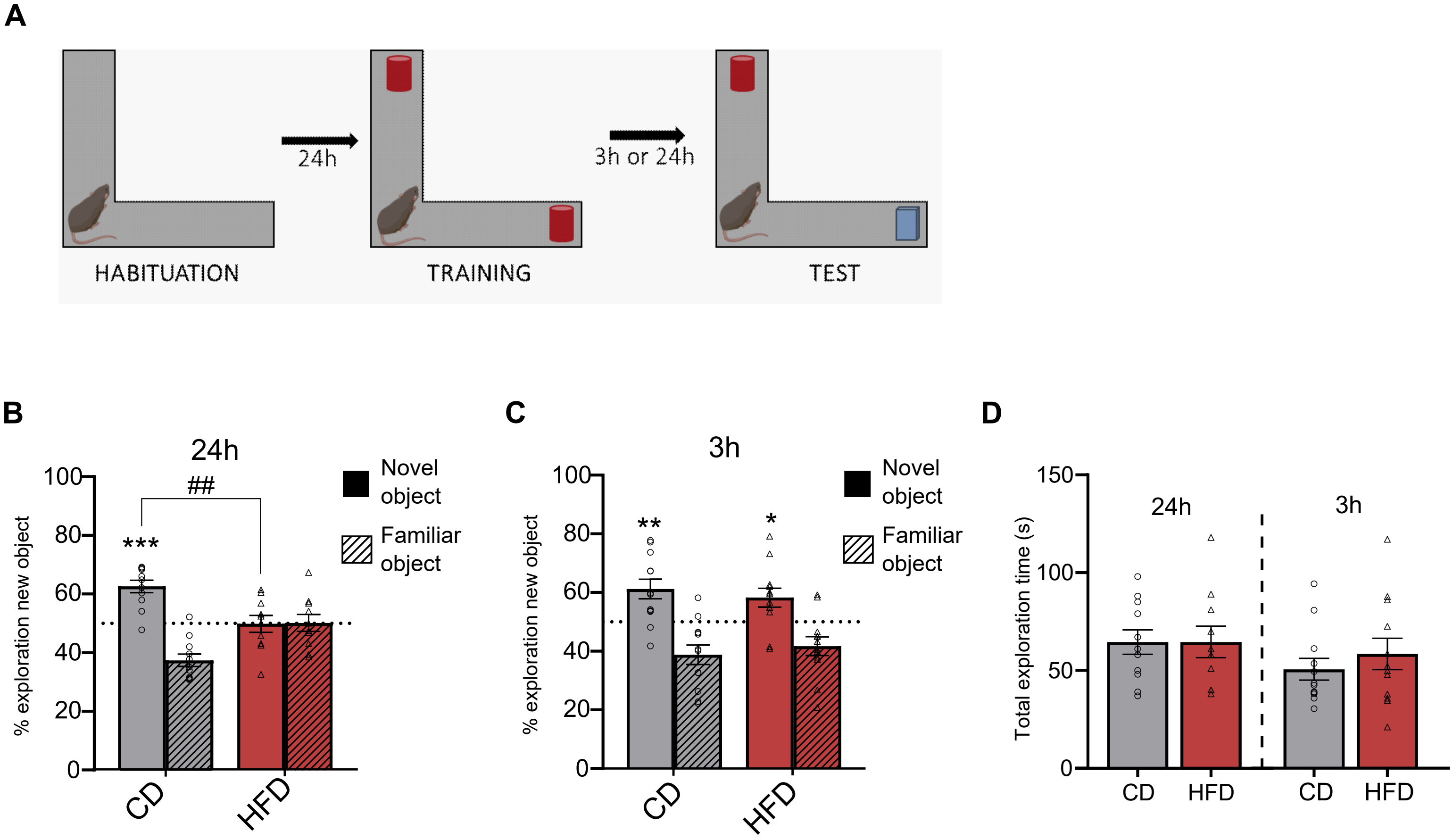

### Supplemental Figure 2

#### Supplementary Figure 2

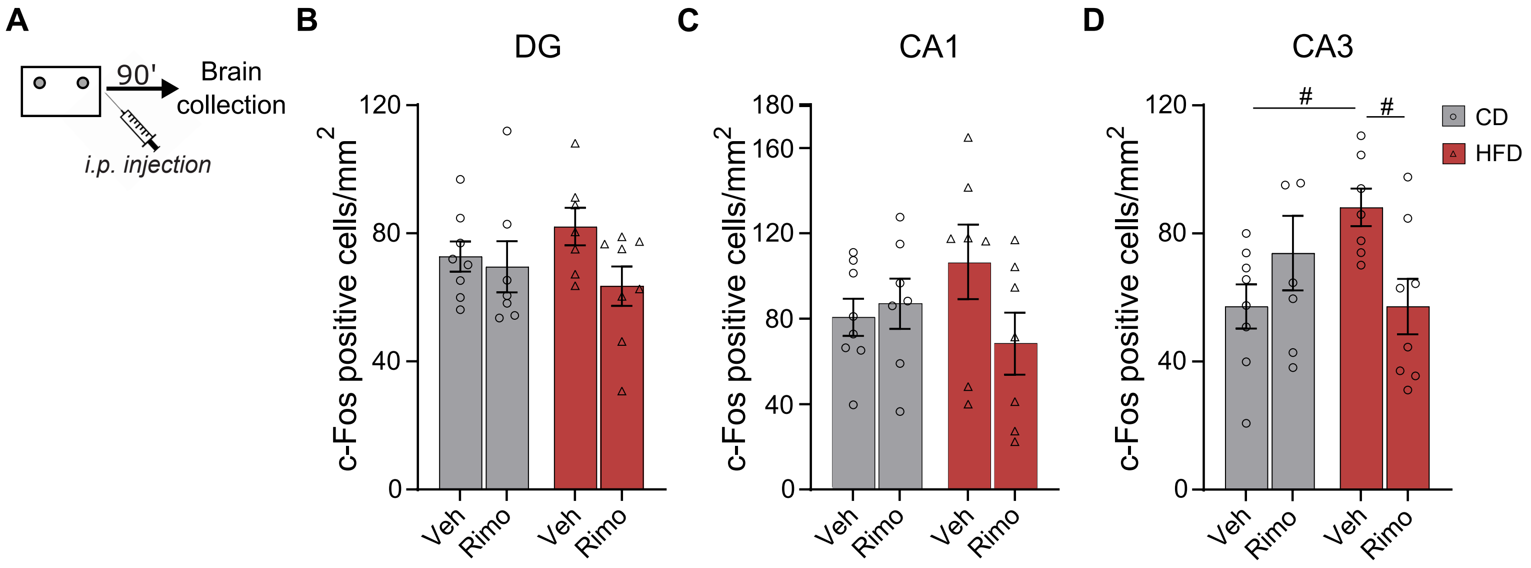

### Supplemental Figure 3

#### Supplementary Figure 3

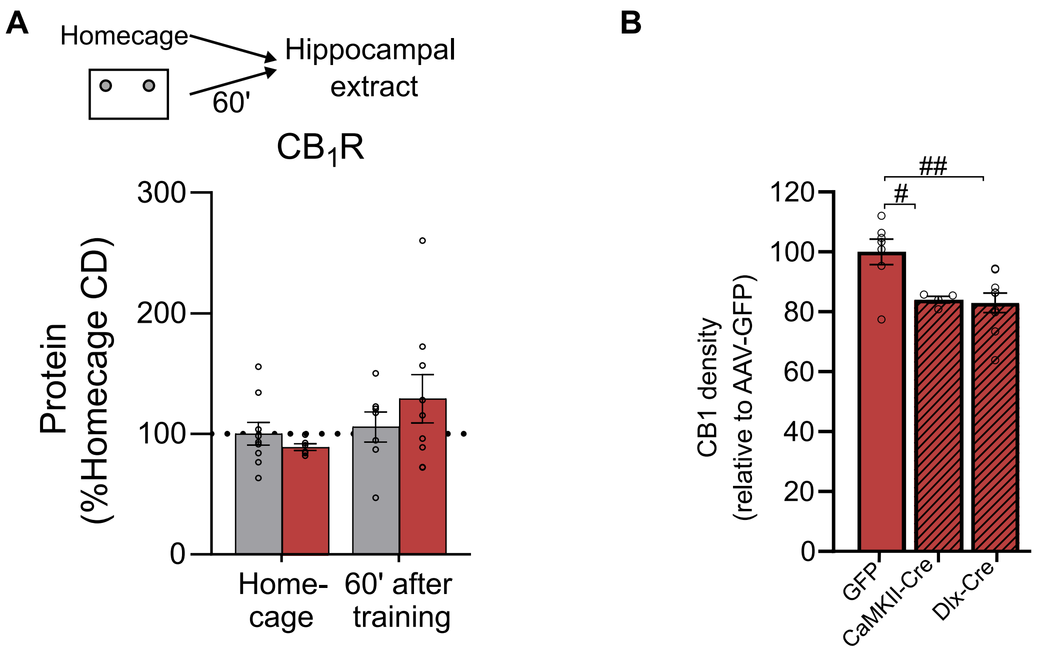

### Supplemental Figure 4

#### Supplementary Figure 4

A

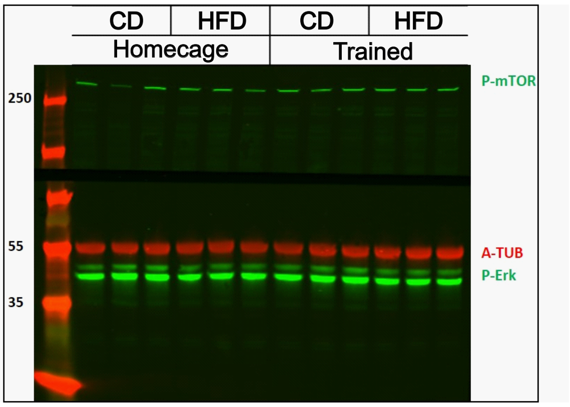


B

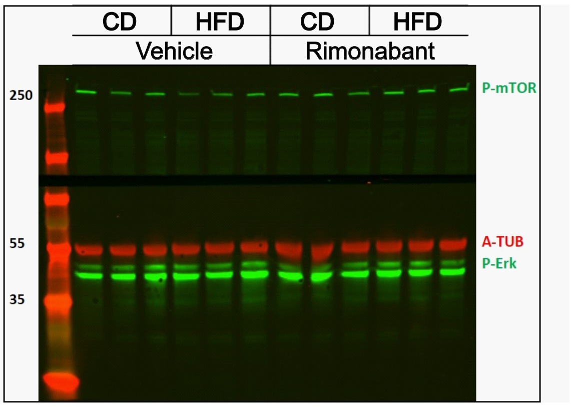
