## Supplemental Table for "Obesogenic diet impairs memory consolidation via the hippocampal endocannabinoid system"

| **Figure 1** | **Experiment** | **Sample size** | **Normality**  (Kolmogorov-Smirnov) | **Analysis** | **t or F-ratios**  *(post-hoc tests reported in Figures)* | **P values** |
| --- | --- | --- | --- | --- | --- | --- |
| 1B | Long-term ORM | 12-15 |  | One sample t-test | CD: t_(14)_= 4.13  HFSD: t_(11)_= 0.16 | 0.029  0.88 |
|  |  |  |  | Unpaired t-test | t_(25)_=2.51 | 0.019 |
| 1C | Short-term ORM | 10-11 |  | One sample t-test | CD: t_(9)_= 5.95  HFSD: t_(10)_= 4.40 | 0.0002  0.0013 |
|  |  |  |  | Unpaired t-test | t_(19)_= 0.76 | 0.45 |
| 1D | Long-term ORM  CB1R antagonist 0h | 11-19 |  | One sample t-test | CD-vehicle: t_(13)_= 3.47  CD-Rim0.5: t_(18)_= 3.03  CD-Rim1: t_(11)_= 0.83  HFSD-vehicle: t_(12)_= 0.08  HFSD-Rim0.5: t_(10)_= 0.27  HFSD-Rim1: t_(13)_= 3.03 | 0.0041  0.0072  0.42  0.94  0.79  0.0097 |
|  |  |  |  | Two-way ANOVA | diet: F_(1,76)_= 3.27  drug: F_(2,76)_= 0.29  diet x drug: F_(2,76)_= 3.70 | 0.074  0.75  0.029 |
| 1E | Long-term ORM  CB1R antagonist 4h | 5-7 |  | One sample t-test | CD-vehicle: t_(4)_= 4.40  HFSD-vehicle: t_(5)_= 0.17  HFSD-Rim1: t_(6)_= 2.08 | 0.012  0.87  0.083 |
|  |  |  |  | One-way ANOVA | F_(2,15)_ = 8.55 | 0.0033 |
| 1F | **Fos expression**  ORM training  CB1R antagonist 0h | 6-8 |  | Two-way ANOVA | *Hippocampus*: diet: F_(1,26)_= 0.32  drug: F_(1,26)_= 2.37  diet x drug: F_(1,26)_= 5.95  *Perirhinal cortex*: diet: F_(1,25)_= 3.71  drug: F_(1,25)_= 0.08  diet x drug: F_(1,25)_= 0.56  *Basolateral amygdala*: diet: F_(1,24)_= 7.56  drug: F_(1,24)_= 0.15  diet x drug: F_(1,24)_= 0.04 | 0.58  0.14  0.029  0.066  0.77  0.46  0.011  0.69  0.84 |
| 1G | **Synaptic plasticity**  Home-cage | 6-9 |  | Two-way repeated measures ANOVA  (post-stimulation) | diet: F_(1,13)_= 0.067  time: F_(19,247)_= 2.75  diet x time: F_(19,247)_= 0.31 | 0.79  0.08  0.99 |
|  |  |  |  | One sample t-test  (1 hour after stim) | CD: t_(8)_= 4.10  HFSD: t_(5)_= 3.19 | 0.0034  0.024 |
|  |  |  |  | Unpaired t-test | t_(13)_= 0.026 | 0.97 |
| 1H | **Synaptic plasticity**  ORM training  CB1R antagonist 0h | 5-10 |  | Three-way repeated measures ANOVA  (post-stimulation) | diet: F_(1,26)_= 0.65  drug: F_(1,26)_= 0.23  time: F_(19,494)_= 2.38  diet x time: F_(19,494)_= 0.63  drug x time: F_(19,494)_= 1.18  diet x drug: F_(1,26)_= 12.08  diet x drug x time: F_(19,494)_= 2.68 | 0.43  0.63  0.093  0.88  0.27  0.0018  0.0002 |
|  |  |  |  | One sample t-test  (1 hour after stim) | CD-vehicle: t_(5)_= 0.20  CD-Rim1: t_(4)_= 5.47  HFSD-vehicle: t_(9)_= 3.79  HFSD-Rim1: t_(4)_= 0.19 | 0.85  0.0054  0.0043  0.85 |
|  |  |  |  | Two-way ANOVA  (1 hour after stim) | diet: F_(1,22)_= 0.41  drug: F_(1,22)_= 0.49  diet x drug: F_(1,22)_= 16.32 | 0.53  0.49  0.0005 |

Supplementary Table 1. Statistical analysis. Related to Figures 1-3 & Supplementary Figure 1.

| **Figure 2** | **Experiment** | **Sample size** | **Normality**  (Kolmogorov-Smirnov) | **Analysis** | **t or F-ratios**  *(post-hoc tests reported in Figures)* | **P values** |
| --- | --- | --- | --- | --- | --- | --- |
| 2A | AEA/2-AG  Home-cage | 21-23 |  | Unpaired t-test | AEA : t_(42)_=0.49  2-AG : t_(42)_=1.07 | 0.62  0.29 |
| 2B | AEA  ORM training | 12-16 |  | Two-way ANOVA | diet: F_(1,78)_= 6.11  time: F_(2,78)_= 93.77  diet x time: F_(2,78)_= = 3.73 | 0.016  0.0001  0.028 |
| 2B | 2-AG  ORM training | 12-16 |  | Two-way ANOVA | diet: F_(1,77)_= 0.09  time: F_(2,77)_= 1.48  diet x time: F_(2,77)_= = 0.056 | 0.76  0.23  0.94 |
| 2C | CB1R mRNA | 5-8 |  | Unpaired t-test | Home-cage: t_(14)_ = 1.17  ORM : t_(9)_ = -2.47 | 0.26  0.035 |
| 2E | Quantification  CB1R decrease | 8-10 |  | Unpaired t-test | Hippocampus : t_(16)_ = 3.55  Basolateral amygdala : t_(16)_ = -0.33 | 0.0038  0.74 |
| 2F | Long-term ORM  Hippocampal  CB1R decrease | 8-15 |  | One sample t-test | CD-GFP : t_(14)_= 2.42  CD-Cre : t_(7)_= 6.25  HFSD-GFP : t_(8)_= 0.49  HFSD-Cre : t_(14)_= 2.26 | 0.029  0.0004  0.64  0.040 |
|  |  |  |  | Two-way ANOVA | diet: F_(1,43)_= 2.90  virus: F_(1,43)_= 4.21  diet x virus: F_(1,43)_= 0.66 | 0.096  0.046  0.42 |
|  |  |  |  |  | *virus effect* (GFP vs Cre)*:* CD  HFD | 0.39  0.045 |
| 2G | Long-term ORM  Hippocampal  CB1R decrease  CaMKII vs DLX | 8-13 |  | One sample t-test | HFSD-GFP : t_(12)_= 0.87  HFSD-CaMKII Cre : t_(8)_= 5.24  HFSD-DLX Cre : t_(7)_= 1.92 | 0.40  0.0008  0.096 |
|  |  |  |  | One-way ANOVA | F_(2,27)_= 5.94 | 0.0073 |

| **Figure 3** | **Experiment** | **Sample size** | **Normality**  (Kolmogorov-Smirnov) | **Analysis** | **t or F-ratios**  *(post-hoc tests reported in Figures)* | **P values** |
| --- | --- | --- | --- | --- | --- | --- |
| 3A | phospho-mTOR | 9-10 |  | Two-way ANOVA | diet: F_(1,35)_= 6.93  training: F_(1,35)_= 0.62  diet x training: F_(1,35)_= 4.54 | 0.012  0.43  0.040 |
| 3A | phospho-ERK | 10-12 |  | Two-way ANOVA | diet: F_(1,40)_= 0.47  drug: F_(1,40)_= 1.10  diet x training: F_(1,40)_= 0.37 | 0.49  0.30  0.55 |
| 3B | phospho-mTOR  CB1R antagonist | 13 |  | Two-way ANOVA | diet: F_(1,48)_= 6.52  drug: F_(1,48)_= 0.64  diet x drug: F_(1,48)_= 1.95 | 0.014  0.43  0.17 |
|  |  |  |  |  | *diet effect* (CD vs HFD): Vehicle  Rim1 | 0.0075  0.41 |
| 3C | Long-term ORM  mTOR inhibitor 0h | 11-12 |  | One sample t-test | CD-vehicle : t_(10)_= 13.54  CD-Temsi: t_(11)_= 13.78  HFSD-vehicle : t_(15)_= 1.87  HFSD-Temsi : t_(18)_= 7.89 | 0.0001  0.0001  0.08  0.0001 |
|  |  |  |  | Two-way ANOVA | diet: F_(1,54)_= 46.79  drug: F_(1,54)_= 7.28  diet x drug: F_(1,54)_= 14.90 | 0.0001  0.0093  0.0003 |
| 3D | Long-term ORM  mTOR inhibitor  4h-8h | 6-8 |  | One sample t-test | HFSD-vehicle 4h: t_(5)_= 2.48  HFSD-Temsi 4h: t_(7)_= 6.45  HFSD-vehicle 8h: t_(6)_= 0.29  HFSD-Temsi 8h: t_(7)_= 0.34 | 0.055  0.0004  0.78  0.75 |
|  |  |  |  | Two-way ANOVA | drug: F_(1,25)_= 4.32  time: F_(1,25)_= 12.02  drug x time: F_(1,25)_= 4.08 | 0.048  0.0019  0.05 |

| **Supplementary**  **Figure 1** | **Experiment** | **Sample size** | **Normality  (Kolmogorov-Smirnov)** | **Analysis** | **t or F-ratios**  *(post-hoc tests reported in Figures)* | **P values** |
| --- | --- | --- | --- | --- | --- | --- |
| 1A | **Fos expression**  ORM training  CB1R antagonist 0h | 6-8 |  | Two-way ANOVA | CA3: diet x drug: F_(1,26)_= 7.84  CA1: diet x drug: F_(1,26)_= 2.84  DG: diet x drug: F_(1,26)_= 1.53 | 0.0095  0.10  0.23 |
